# Tissue Fluidity Promotes Epithelial Wound Healing

**DOI:** 10.1101/433557

**Authors:** Robert J. Tetley, Michael F. Staddon, Shiladitya Banerjee, Yanlan Mao

## Abstract

Epithelial tissues are inevitably damaged from time to time and must therefore have robust repair mechanisms. The behaviour of tissues depends on their mechanical properties and those of the surrounding environment^1^. However, it remains poorly understood how tissue mechanics regulates wound healing, particularly in *in vivo* animal tissues. Here we show that by tuning epithelial cell junctional tension, we can alter the rate of wound healing. We observe cells moving past each other at the wound edge by intercalating, like molecules in a fluid, resulting in seamless wound closure. Using a computational model, we counterintuitively predict that an increase in tissue fluidity, via a reduction in junctional tension, can accelerate the rate of wound healing. This is contrary to previous evidence that actomyosin tensile structures are important for wound closure^2–6^. When we experimentally reduce tissue tension, cells intercalate faster and wounds close in less time. The role we describe for tissue fluidity in wound healing, in addition to its known roles in developing^7,8^ and mature tissues^9^, reinforces the importance of the fluid state of a tissue.

## Main Text

To investigate the role of tissue mechanics in wound healing, we studied tissue repair in the epithelium of *ex vivo Drosophila* wing imaginal discs by live time-lapse imaging. After wounding wing discs by laser ablation, an actomyosin purse string assembles at the wound’s leading edge (Fig. 1a, Supplementary Video 1), as in other systems^5,10–13^. To understand the repair process in more detail, we quantitatively analysed^14^ wound morphology over the time course of wound closure. We observe three distinct phases after wounding: recoil, fast closure and slow closure (Fig. 1b, Supplementary Video 2). Immediately after wounding, the wound area increases, due to a release of tissue tension by the ablation. The wound area then reduces in time, with an initial fast phase. However, after reaching approximately 50% of the original wound area, the rate of wound closure decreases dramatically, until closure.

**Figure 1.**
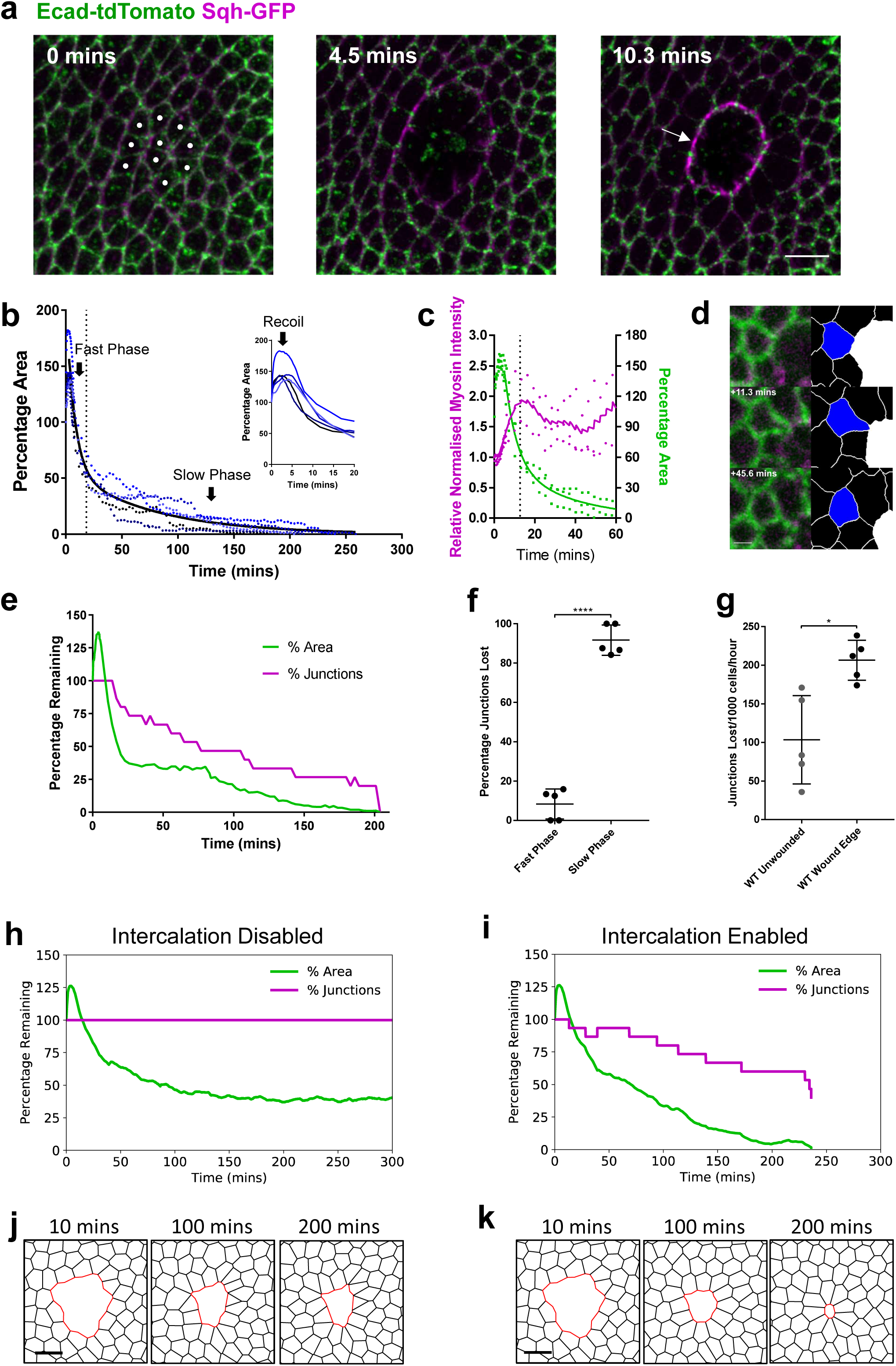
Wing disc wound closure is punctuated by wound edge intercalation, which can drive wound closure. **a**, Early stages of wound closure in a *sqh^AX^*^3^; *sqh*-GFP, Ecad-tdTomato wing imaginal disc. Cell outlines are marked by Ecad-tdTomato (green) and Myosin II by Sqh-GFP (magenta). Cells which will be ablated are marked by white circles at 0 mins. Within the first 10 minutes after wounding, a strong accumulation of Myosin II can be seen at the wound’s edge in the manner of a purse string (arrow). Images are maximum intensity projections of deconvolved image stacks. Scale bar = 5μm. **b**, Dynamics of wing disc wound closure. Percentage of original wound area is plotted over time for 5 WT wing discs expressing Ecad-GFP (blue dots, the same 5 wing discs are used for all subsequent WT analysis unless otherwise stated). Inset: first 20 mins, showing early expansion (recoil) of the wound. A two-phase exponential decay curve (black line) is fitted to the data after 3 minutes, when the wound begins to reduce in area until close. The transition between fast and slow closure phases of the two-phase exponential decay is marked by a dotted line (18.37 mins). **c**, Quantification of Myosin II purse string intensity (magenta, left y-axis) and wound percentage area (green, right y-axis) for 3 *sqh^AX3^*; sqh-GFP, Ecad-tdTomato wing discs during the first hour of wound closure. A two-phase exponential decay curve has been fitted to the area data (green line) and a LOWESS (smoothing window of 10) curve to the Myosin II intensity data (magenta line). The transition between fast and slow closure phases is shown with a dotted line (12.71 mins) **d**, Example of a single wound edge intercalation in an Ecad-GFP; *sqh*-mCherry, *pnr*-GAL4 wing disc. Raw maximum intensity projection (left) and skeletonised images (right, intercalating cell in blue, wound in white) are shown. The junction shared between the intercalating cell and the wound shrinks to a point and a new junction grows in the orthogonal direction. Scale bar = 3μm. **e**, Quantification of the percentage of starting wound edge junctions (magenta) and wound percentage area (green) for a single Ecad-GFP wing disc wound. The percentage of junctions remaining on the wound’s edge reduces as intercalations occur until the wound fully closes. **f**, Quantification of the percentage of junctions lost during the fast (early) and slow (late) phases of WT wound closure. Significantly more junctions are lost during the slow phase (unpaired *t*-test, n=5, *t*=17.13, *df*=*8*, *p*<0.0001). Error bars = SD. **g**, Quantification of intercalation rate in unwounded WT tissues and at WT wound edges. The intercalation rate is significantly higher at the wound edge (unpaired *t*-test with Welch’s correction, n=5, *t*=3.667, *df*=5.567, *p*=0.012). Error bars = SD. **h, i**, Vertex model simulations. Percentage of initial wound area and wound junctions after ablation with intercalations (**h**) disabled, and (**i**) enabled. **j, k**, Vertex model simulation images after ablation with intercalations (**j**) disabled, and (**k**) enabled.

We hypothesised that myosin II (MyoII) dynamics in the purse string might regulate changes in wound closure rate and quantified the intensity of MyoII in the purse string during the closure process (Fig. 1c). MyoII intensity increases after wounding, before peaking at roughly twice the initial intensity. This peak coincides with the transition between fast and slow closure phases. The saturation of MyoII before complete wound closure suggested that other cell behaviours might be required to generate forces to complete wound closure. By closely examining cell behaviours around the wound, we observe that cells at the wound edge readily undergo intercalation (Fig. 1d, e, Supplementary Video 2). This cell behaviour differs to those associated with wound healing in other *Drosophila* tissues, such as cell fusion^15,16^, polyploidisation^16^ and cell intercalation further from the wound edge^17^. During wound edge intercalations, junctions in contact with the wound shrink to a single vertex and new junctions grow in the orthogonal direction (Fig. 1d). As a result, the number of cells in contact with the wound decreases over time (Fig. 1e, Supplementary Fig. 1a-d). The rate at which wound edge cells intercalate is roughly double that of an unwounded tissue (Fig. 1g), while the majority of intercalation events occur during the slow phase of wound closure (Fig. 1f), indicating that wound edge cell intercalation might be required to promote the completion of wound closure.

To quantitatively test the role of wound edge cell intercalation, we developed a computational vertex model^18^ for wound closure (Supplementary Fig. 2, see methods). We parameterised the model so that edges contacting the wound gradually increase in tension compared to the surrounding tissue, to mimic the assembly of the contractile actomyosin purse string. To capture experimentally observed fluctuations in junctional MyoII^7^, we introduced fluctuations in line tension. Without introducing intercalation events into the model, simulated wounds are unable to close (Figs. 1h, j, Supplementary Video 3). By contrast, when intercalation is enabled in the model (see methods), wounds are able to close (Figs. 1i, k, Supplementary Video 4), supporting the hypothesis that intercalation at the wound edge is necessary for wing disc wound closure.

The vertex model predicts that in the absence of intercalation, cells around the wound become increasingly elongated towards the centre of the wound (Fig. 2a), unlike in simulations with intercalations enabled (Fig. 2b). This led us to hypothesise that wound edge intercalation has a crucial role in maintaining cell shape and tissue patterning. Indeed, wing disc cells appear regularly packed immediately after wound closure (Fig. 2c) and the polygon distribution of wound edge cells is restored upon healing (Fig. 2d). To test our vertex model’s prediction that intercalation preserves cell shape, we quantified cell elongation in the first three rows of cells away from the wound in wing discs (Fig. 2e, Supplementary Video 5). Whereas cells in the second and third rows undergo little change in elongation during wound closure (Fig. 2f), cells in the first row behave differently and undergo a transient increase in elongation, before returning to their original shapes prior to wound closure (Figs. 2f-h). Cells return to their original shape during the slow phase of closure (Fig. 2f), when the majority of intercalations occur, supporting the role of intercalation in preserving cell shape.

**Figure 2.**
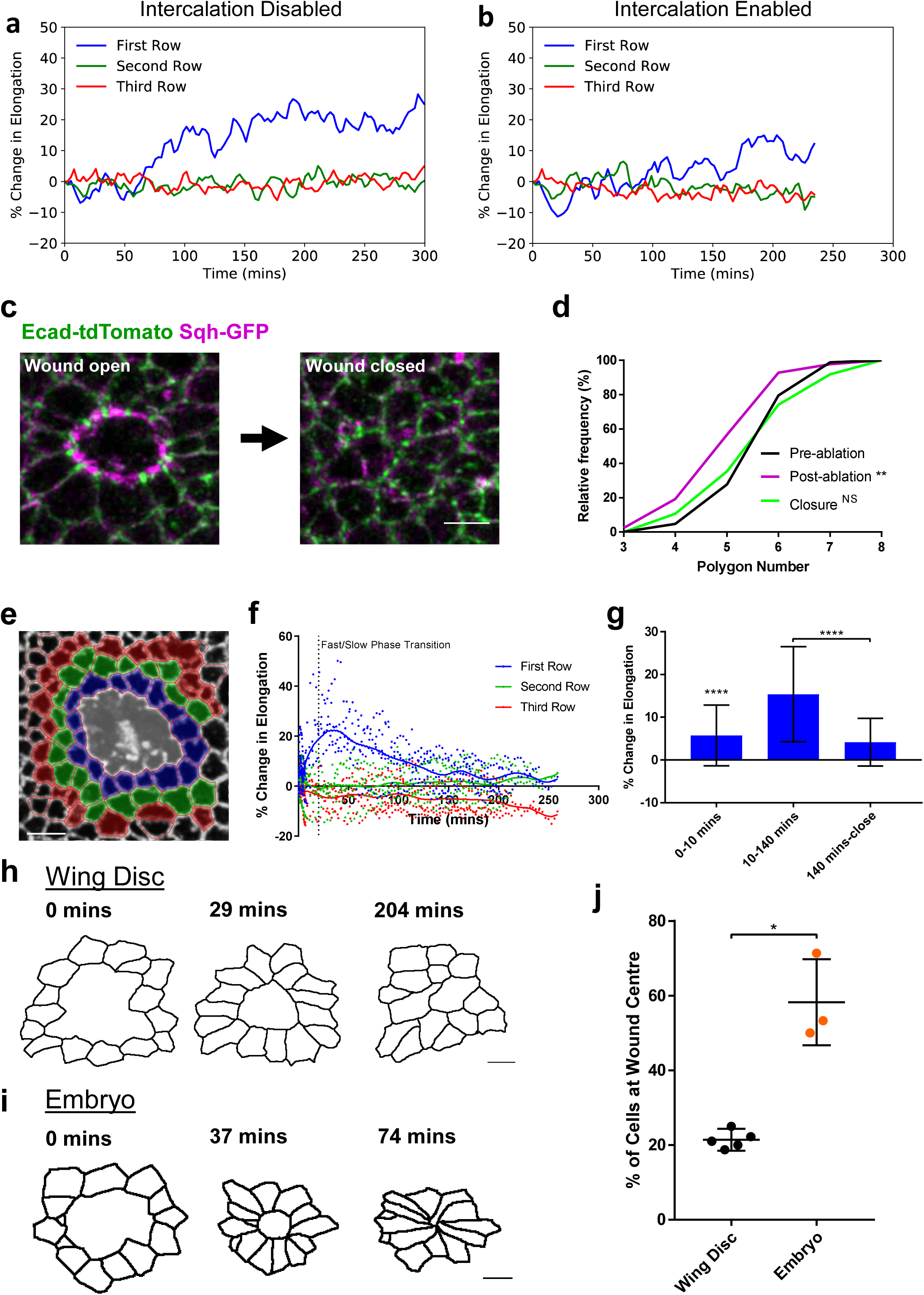
Wound edge intercalation preserves cell shape. **a**, **b**, Vertex model simulations. Percentage change in cell elongation, calculated by dividing the major axis by the minor axis of an ellipse fit to each cell, for the first three rows of cells around the wound with intercalations (**a**) disabled, and (**b**) enabled. **c**, Maximum intensity projection images of a wound in a *sqh^AX3^*; *sqh*-GFP, Ecad-tdTomato wing disc before (left) and immediately after (right) wound closure (scale bar = 3μm). **d**, Quantification of wound edge cell polygon number before ablation, immediately after ablation and immediately after wound closure. The distribution of polygon number is significantly shifted left after ablation (Kolmogorov-Smirnov Test, n=5, D=0.2892, *p*=0.0019) but is restored upon closure (Kolmogorov-Smirnov Test, D=0.07583, p=0.9692). **e**, Colour coding of the first three rows away from the wound edge (first row blue, second row green, third row red). Image is an adaptive projection of Ecad-GFP overlaid by skeletonised cell outlines. Scale bar = 5μm. **f**, Quantification of the percentage change in mean cell elongation for the first three rows of cells (colour coding as in e) over time for 5 WT wounds. LOWESS regression curves (smoothing window = 10) are shown. The transition between fast and slow closure phases is marked by a dotted line (18.37 mins). Elongation transiently increases for just the first row of cells. **g**, Percentage change in mean elongation of first row cells for three time windows; 0–10 mins (wound recoil), 10–140 mins (early wound closure), 140 mins – close (late wound closure). Data is pooled from the data set in (f). Cells were significantly elongated during the recoil phase (Wilcoxon Signed Rank Test, *p*<0.0001). Cell were significantly more elongated during early wound closure than during late wound closure (Kolomogorov-Smirnov Test, D=0.5703, *p*<0.0001). Error bars = SD. **h, i**, Skeletonised cell outlines of cells starting at the wound edge at three time points in a (**h**) single WT wing disc (scale bar = 3μm) and a (**i**) single WT stage 13 embryo (scale bar = 5μm). **j**, Quantification of the percentage of cells remaining close to the wound’s centre after closure, as a measure of intercalation. A significantly higher percentage of cells remain close to the wound centre in embryos (n=3) compared to wing discs (n=5) (unpaired *t*-test with Welch’s correction, *t*=5.466, *df*=2.103, *p*=0.0285).

These findings suggested that intercalation events help maintain cell shape and that wound healing in an epithelium that does not intercalate will lead to cell deformation. To test this idea further, we compared wound edge intercalation in the wing disc to the *Drosophila* embryonic ectoderm, an epithelium in which cells do not return to their original shape (Fig. 2i). Following wound closure in the *Drosophila* embryonic ectoderm, cells can be up to twice as elongated as they were prior to wounding^17^. Supporting the model’s predictions, there are significantly fewer wound edge intercalations in embryos prior to wound closure (Fig. 2j).

These results suggested that wound closure is controlled by two dynamic properties of the wounded tissue: the rate of wound edge intercalation and the tension in the purse string. In support of this view, simulations using our vertex model demonstrate that rates of cell division (Supplementary Fig. 3e, f) and magnitudes of edge tension fluctuations (Supplementary Fig. 3c, d) have little effect on wound closure rate. We therefore used the model to test the relative roles of intercalation rate and purse string tension in wounded tissues. We found that the rate of intercalation in the tissue can be quantitatively tuned by modulating the line tension term, which represents cell-cell interfacial tension in our model^7^ (Fig. 3a). Intercalation rate can also be tuned by modulating the perimeter contractility (Supplementary Fig. 3a, b), however, as there is redundancy in the 2 terms in representing MyoII contraction, we chose to only vary the line tension term in subsequent simulations. As cell line tension decreases, the rate of intercalation in the tissue increases, which we term an increase in “tissue fluidity” – the rearrangement of cells relative to each other being analogous to molecules in a liquid. Increasing either purse string tension or tissue fluidity (by decreasing line tension) reduces the time taken for wounds to close (Fig. 3b). Although reducing line tension throughout the system increases bulk tissue fluidity (Fig. 3c), the effect is more enhanced at the wound edge (Fig. 3d). Unexpectedly, we find a region of parameter space where a reduction in purse string tension is more than compensated for by an increase in tissue fluidity, leading to accelerated wound closure (Figs. 3b, e-f, magenta). This suggests that tissue fluidity can act as a predominant driver of wound closure and that the purse string may only be providing a directional cue at late closure stages, despite providing the initial driving force.

**Figure 3.**
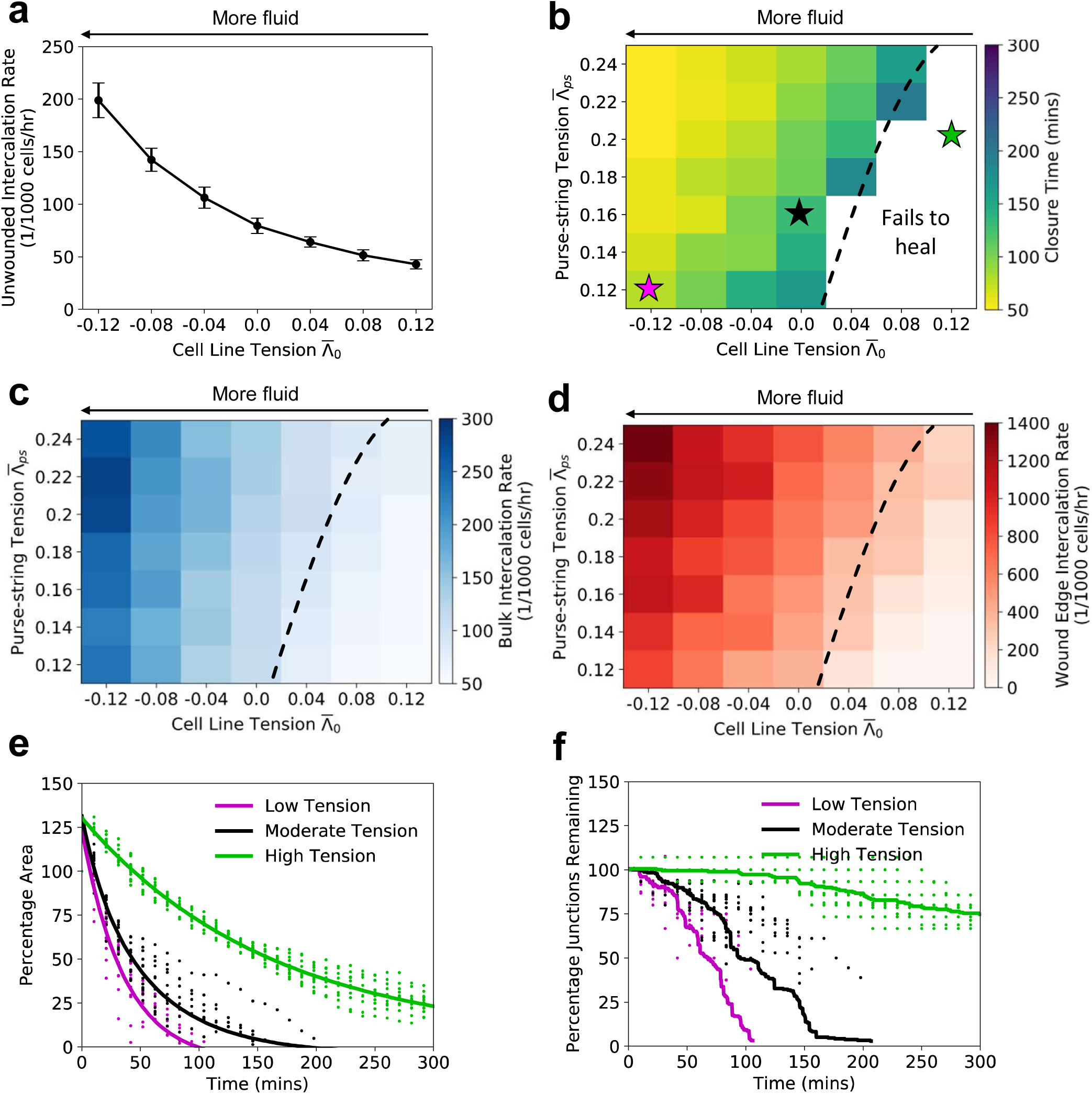
Reducing tissue contractility enhances fluidity and can speed wound closure. All figures are vertex model simulations. **a**, Intercalation rate in an unwounded tissue against mean cell line tension. Error bars = SD. **b**, Mean wound closure time against cell line tension and purse-string tension. The white region indicates parameter space where wounds fail to close within 300 minutes. The dashed line represents a transition between healing and non-healing regions. The colored stars indicate the parameters used in (e) and (f). **c**, Mean bulk intercalation rate in wounded tissues, and **d**, mean wound edge intercalation rate against cell line tension and purse-string tension. **e**, Percentage of initial area over time for low, moderate, and high tension cases. Points are from simulations, and lines are fit dual exponential curves. **f**, Percentage of initial wound junctions over time for low, moderate, and high tension cases. Points are from simulations; lines are the average over all simulations. For all combinations of tension, n = 12 simulations for each.

To test these predictions from our vertex model further, we sought to experimentally perturb cell edge tension. In epithelial tissues, cell edge tension is governed by the activity of non-muscle MyoII (Fig. 4a). To test the roles of purse string tension and tissue fluidity in the wing disc, we genetically modulated the activity of MyoII in the wing pouch epithelium. To increase tension, we performed RNAi against the *Myosin binding subunit (Mbs)* of the Myosin Phosphatase, a phosphatase that inactivates MyoII by dephosphorylating its regulatory light chain^19^ (Spaghetti squash (Sqh) in *Drosophila)*. To decrease tension, we performed RNAi against *Rho-kinase (Rok)*, a kinase that activates MyoII by phosphorylating Sqh^20,21^. We confirmed the effect of these genetic perturbations on tension by quantifying vertex recoil rates (a greater recoil rate implying higher tension) after single junction ablations (Supplementary Fig. 5). We then wounded these wing discs (Supplementary Fig. 4a-d) and compared the dynamics of wound closure to wildtype (WT) wing discs. In *Mbs* RNAi wing discs, where tension is high, wounds fail to close within the imaging time window (Figs. 4b, c, Supplementary Fig. 4a, Supplementary Fig. 6, Supplementary Video 6). Furthermore, wound edge cell intercalation is almost entirely abolished (Figs. 4c, e-f, Supplementary Fig. 6, Supplementary Video 6), supporting the importance of intercalation in promoting wound closure.

**Figure 4.**
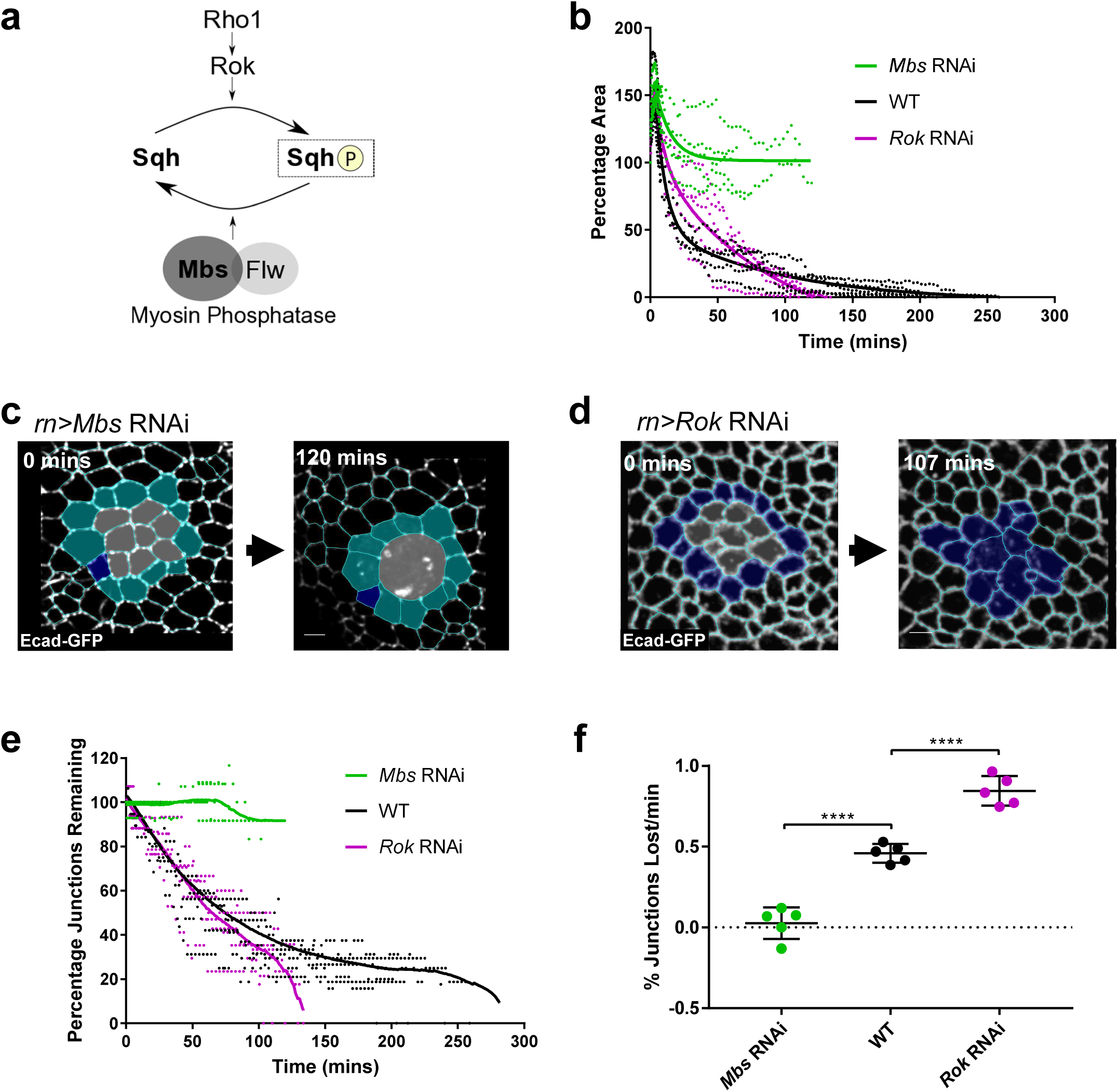
Myosin activity controls tissue fluidity and wound closure rate. **a**, Activation of myosin II by phosphorylation of its regulatory light chain (Sqh) can be performed by Rho kinase (Rok) downstream of Rho1. Myosin II inactivation by Sqh dephosphorylation can be performed by the Myosin Phosphatase comprising the Myosin binding subunit (Mbs) and catalytic subunit (Flapwing, Flw). **b**, Quantification of wound closure (as percentage of start wound area) over time in *Mbs* RNAi (green, n=5) and *Rok* RNAi (magenta, n=5) wing discs, compared to WT wound closure (black, n=5). Two-phase exponential decays are fitted after 3 minutes. *Rok* RNAi wounds close faster than WT and *Mbs* RNAi wounds fail to close. **c, d**, Examples of wound healing in (**c**) *Mbs* RNAi and (**d**) *Rok* RNAi before wounding (left) and after wound closure (*Rok* RNAi) or when further segmentation become impossible (*Mbs* RNAi). Cells are colour coded according to whether they undergo intercalation (dark blue) or not (cyan). Images are adaptive projections of Ecad-GFP overlaid by skeletonised cell outlines (scale bars = 3μm). **e**, Quantification of the percentage of initial wound edge junctions over time for *Mbs* RNAi and *Rok* RNAi wing discs (colours and n numbers as in b). LOWESS curves (smoothing window of 5) are fitted to the data. Junctions are lost from the wound edge through intercalation faster in *Rok* RNAi wounds than in WT. *Mbs* RNAi wounds lose very few junctions. **f**, Quantification of mean intercalation rate for *Mbs* RNAi, WT and *Rok* RNAi wounds (colours and n numbers as in b). Intercalation rate is significantly higher in *Rok* RNAi wounds (unpaired *t*-test, *t*=8.026, *df*=8, *p*<0.0001) and significantly lower in *Mbs* RNAi wounds (unpaired *t*-test, *t*=8.503, *df*=8, *p*<0.0001) compared to WT. Error bars = SD.

In *Rok* RNAi wing discs (Supplementary Fig. 7, Supplementary Video 7), in which tension is reduced, wounds close faster than in WT wing discs. While these wounds initially close more slowly than WT wounds, they eventually overtake them and unexpectedly close in roughly half the time of WT wounds (Figs. 4b, d, Supplementary Fig. 4a). This is accompanied by an increase in the rate of wound edge intercalation (Figs. 4d-f). One consequence of decreasing MyoII activity through *Rok* RNAi, is that it is likely to increase tissue fluidity, while simultaneously decreasing purse string tension^3,22^. The observation that the *Rok* RNAi wounds close faster despite a weakened purse string is in agreement with the regions of parameter space in our vertex modelling, where a weakened purse string can be overcome by an increase in tissue fluidity (Figs. 3b, e-f, magenta).

We have demonstrated that wing disc wound closure is dependent on cell-cell intercalation-driven tissue fluidity. Intercalation a few rows of cells away from the wound is also known to be important for wound closure in the *Drosophila* embryonic ectoderm^17^. However, the embryonic intercalations resemble those characteristic of *Drosophila* germband extension^23^, relying on active polarised flows of actomyosin^17^, while in the wing disc MyoII activity is inhibitory to intercalation and fluidity. Polarised actomyosin activity may be a hallmark of healing in more naïve embryonic tissues and global junctional tension-based fluidity may dominate in more mature tissues. Observations in human bronchial epithelial cell (HBEC) layers support this hypothesis. HBEC layers demonstrate a progressive loss of fluidity after becoming confluent^9^, likely sharing parallels with wound healing, as in both situations the epithelium must explore multiple cellular conformations before reaching a new homeostatic state.

Changes in tension-based fluidity, such as those we have induced by MyoII perturbation, can be considered as jamming/unjamming transitions^24,25^. Cell layers can transition between behaving like a fluid (unjammed) with many rearrangements and a solid (jammed), lacking the ability to rearrange^24,25^. Modelling has described how jamming/unjamming transitions can occur due to changes in junctional tension and cell-cell adhesion^9^. These two properties combine to control the magnitude of a mechanical energy barrier, which must be overcome for cells to rearrange relative to each other^9,24^, allowing an epithelium to behave as a fluid. By reducing tension in the wing disc, we are likely lowering this energy barrier, allowing cells to rearrange more. Simulations demonstrate that, with intercalations, tissues can transit to a lower energy state, which may allow further intercalations to occur (Supplementary Fig. 8a-c).

The dynamics of wound closure in wing discs can be explained purely through junctional dynamics in our vertex model, rather than previously described cell-crawling based migration^26–29^. Closing a wound by junctional dynamics alone may be a mechanism through which epithelial integrity and function can be maintained. Furthermore, our vertex model simulations suggest that wounds can close simply by introducing a purse string, without the need to actively change any chemical or mechanical properties of the surrounding tissue, provided its original state is permissive to intercalations. The increased fluidity we observe at the wound edge in wing discs is therefore an emergent property defined by the pre-existing junctional tension of the tissue in response to an actomyosin purse string. Changing tension of a wounded tissue, as we have done in the wing disc, therefore presents an attractive therapeutic target that could be used to accelerate wound healing in the future.

## Acknowledgements

Thank you to all members of the Mao group, Martin Raff, David Ish-Horowicz and Michael Murrell for providing feedback on the manuscript. Thank you to Andreas Hoppe and Davide Heller for developing additional plugins in Epitools for this work.

## Author Contributions

RJT and YM conceived the project. RJT performed the experiments and analysed the data. MFS and SB developed the vertex model of wound healing. MFS ran simulations and analysed the data. RJT, MFS, SB and YM wrote the manuscript.

## Author Information

RJT is funded by a Medical Research Council Skills Development Fellowship (MR/N014529/1). MFS is supported by an EPSRC funded PhD Studentship at the UCL Department of Physics and Astronomy. SB acknowledges support from a Strategic Fellowship at the UCL Institute for the Physics of Living Systems. YM is funded by a Medical Research Council Fellowship (MR/L009056/1), a UCL Excellence Fellowship, and a NSFC International Young Scientist Fellowship (31650110472). This work was also supported by MRC funding to the MRC LMCB University Unit at UCL (award code MC_U12266B).

## Methods

### *Drosophila* strains

*Drosophila* stocks were raised on conventional cornmeal media at 25°C. The following alleles and transgenes were used experimentally: *shg*-GFP^1^ (II, referred to as Ecad-GFP), *sqh*-mCherry^2^ (III), *rn*-GAL4 (III, MiMIC insertion), UAS-R*ok*-RNAi (II, VDRC KK Library), UAS-*Mbs*-RNAi (II, VDRC KK Library), *sqh*-GFP^3^ (II), *shg*-tdTomato^1^ (II, referred to as Ecad-tdTomato), *sqh^AX3^* (X)^4^. The experimental genotypes used and their corresponding data are shown in Table 1.

**Table 1.** *Drosophila* experimental genotypes and corresponding data.

| Genotype | Corresponding Figures and Videos |
| --- | --- |
| *sqh*<sup>AX3</sup>; *sqh*-GFP, Ecad-tdTomato | Fig. 1a, c, Fig. 2c, Supplementary Video 1 |
| *yw*; Ecad-GFP/+; *rn*-GAL4/+ (referred to as WT) | Fig. 1b, e-g, Fig. 2 d-h, j, Fig. 4b, e-f, Supplementary Fig. 1a-d, Supplementary Fig. 4a-d, Supplementary Fig. 5, Supplementary Video 2, Supplementary Video 5 |
| Ecad-GFP; *sqh*-mCherry, *pnr*-GAL4 | Fig. 1d |
| *yw*; Ecad-tdTomato | Fig. 2i, j |
| *yw*; Ecad-GFP/UAS-*Mbs*-RNAi; *rn*-GAL4/+ (referred to as *Mbs*-RNAi) | Fig. 4b-c, e-f, Supplementary Fig. 4a-d, Supplementary Fig. 5, Supplementary Fig. 6a-e, Supplementary Video 6 |
| *yw*; *Ecad-GFP/UAS-Rok-RNAi*; *rn-GAL4/+* (referred to as *Rok-RNAi*) | Fig. 4b, d-f, Supplementary Fig. 4a-d, Supplementary Fig. 5, Supplementary Fig. 7a-e, Supplementary Video 7 |

### Live imaging of wing imaginal discs

Late third instar wing imaginal discs were cultured in Shields and Sang M3 media (Sigma) supplemented with 2% FBS (Sigma), 1% pen/strep (Gibco), 3ng/ml ecdysone (Sigma) and 2ng/ml insulin (Sigma). Wing discs were cultured under filters, as described elsewhere^5^. Wing discs were imaged on a Zeiss LSM 880 microscope with Airyscan at 512×512 resolution with a 63x objective (NA 1.4) at 5x zoom. Laser power of 0.2–0.3% was used and images were captured with a 0.5μm z-spacing. Time intervals varied depending on the experiment. For single channel wounding experiments, the first 25 images were captured with no time interval (to visualise the fast, early dynamics of wound closure, using a sufficient z-stack depth to include all cells in the field of view). For two channel wounding experiments, the first 12 images were captured with no time interval. Subsequent imaging was then performed with a time interval of 3 minutes for both single and two channel experiments, with a total z-stack depth of approximately 40 μm, until wounds closed. For experiments measuring unwounded intercalation rates, a time interval of 3 minutes was used for a total imaging time of 2 hours.

### Wounding of tissues

Wing discs were wounded using a pulsed Chameleon Vision II TiSa laser (Coherent), tuned to 760nm at 45% power. Ablation was performed on small, manually defined circular regions of interest (ROIs) that coincided with the tricellular junctions shared between all cells to be ablated. This was necessary, as larger regions of interest produced significant autofluorescent scarring that made subsequent image analysis impossible. Furthermore, ablation of larger regions led to cavitation, which masked the initial recoil dynamics of the wound. Ablation was performed in a single z-plane, at the level of the adherens junctions.

### Segmentation and tracking of wound time-lapse images

Ecad time-lapse images were first deconvolved using Huygens (Scientific Volume Imaging). Deconvolved images were segmented, tracked and analysed using Epitools^6^. The following MATLAB-based analysis modules were implemented in the following order: Adaptive projection (manually corrected with an additional plugin, Andreas Hoppe, unpublished), Contrast Enhancement (CLAHE), Cell segmentation, Automatic seed tracking, Resegmentation, Generate skeletons. The settings used for “Adapative projection” and “Cell segmentation” were selected on a case-by-case basis, to give the best result for each time-lapse image. Default settings were used for “Contrast Enhancement (CLAHE). “Automatic seed tracking” was implemented so that each cell had a single seed, where possible. Corrected seeds were used for “Re-segmentation” and the resulting corrected segmentation was used to “Generate skeletons”. The remaining segmentation and tracking was performed using Epitools’ Icy Plugins. Skeletons were manually corrected using “CellEditor”. The corrected skeletons were used to implement “CellGraph”, which was run using the “STABLE_MARRIAGE” algorithm for tracking with a Propagation Limit of 5 frames without cutting any border lines. Any tracking errors were manually corrected (additional tool, Davide Heller, unpublished).

### Quantitative analysis of wound time-lapse images

The following data was quantified and exported using the “CellOverlay” Icy plugin in Epitools^6^. Statistical analysis and curve fitting was performed in Prism (GraphPad): *Wound area* – the killed cells and resulting wound were manually selected using the “CELL_COLOR_TAG” tool and data exported. Wound area was then expressed as a percentage of the total area of all the killed cells.

#### Wound edge junctions remaining

the initial number of wound edge cells was counted manually prior to wounding. The polygon number of the wound was quantified using the “CELL_COLOR_TAG” tool. The wound polygon number was expressed as a percentage of the initial number of wound edge cells. When percentage of junctions remaining data was pooled from multiple wing discs a LOWESS curve was fitted to the data.

#### Distinction of fast and slow closure phases

a two-phase exponential decay curve was fitted to the cell area data pooled from multiple wing discs. The fast and slow phases of closure are represented by the ranges of the fast and slow exponential decays respectively.

#### Intercalation rates

Intercalation rates were calculated in two ways: (1) To calculate the mean wound edge intercalation rate for an entire time-lapse image, the total percentage of junctions lost was divided by the total time to give a value of percentage of junctions lost per min (Fig. 4f). (2) To calculate the intercalation rate for unwounded tissues the “EDGE_T1_TRANSITIONS” tool was run and data exported. Analysing the raw data, any T1 transition that was maintained for >1 time point was scored as a junction loss event (to allow comparison to wound edge junction losses during wound edge intercalation). The total number of cells was quantified by exporting data from the “TRACKING_STABLE_ONLY” tool. Intercalation rate was calculated as the number of junctions lost per 1000 cells per hour. The same units were used for wound edge intercalations. However, because a junction loss during a T1 transition in an unwounded tissue involves 4 cells and a junction loss at the wound edge involves only 3 cells, the wound edge intercalation rate was adjusted by a factor of 0.75 (Fig. 1g).

#### Polygon distributions of wound edge cells

initial wound edge cells were selected manually using the “CELL_COLOR_TAG” tool, prior to wounding. These identities were propagated through all time points. Data was exported and the polygon distributions quantified by pooling all data from all wound edge cells in all wing discs. Polygon distributions were quantified at time points prior to and immediately after wounding and immediately after closure.

#### Cell elongation

cell elongation was calculated by dividing the major and minor axis length of best fit ellipses (exported from the “CELL_COLOR_TAG” tool). The mean elongation of pooled cells was calculated prior to wounding. Changes in mean cell elongation were expressed as a percentage of the initial mean elongation.

Overlay images were generated by selecting the relevant “CellOverlay” tool layer.

### Myosin II quantification

Myosin II was quantified during the first hour post wounding of *sqh^AX3^*; *sqh*-GFP, Ecad-tdTomato wing discs, using Sqh-GFP as a reporter. Because these experiments were performed in a *sqh^AX3^* null background, all molecules of Myosin II were tagged with GFP. First, raw (not deconvolved) Sqh-GFP images were background subtracted using the Rolling Ball tool in FIJI with a radius of 12 pixels. Maximum intensity projections (MIPs) of Sqh-GFP images were then generated, which excluded any signal from the overlying peripodial membrane cells. The CellGraph function of Epitools was run on the Sqh-GFP MIPs using the segmented skeleton images from the corresponding deconvolved Ecad-tdTomato channel (segmentation performed as above).

Junctions were selected manually using the “EDGE_COLOR_TAG” CellOverlay tool in Epitools^6^. Edge lntensity Buffer and Vertex Intensity Buffer were set to 3 and Selection Mode 1 was used to exclude vertices from the quantification. Two groups of junctions were selected in each movie; 1) junctions in contact with the wound edge in each frame and 2) 10 junctions in the surrounding tissue that persisted for the entire time window of quantification, were more than one cell diameter away from the wound and were not associated with a cell division event. Mean Sqh-GFP intensities were exported for all tagged junctions, with the mean calculated from the top 90% of pixel intensities for each junction (to exclude any dark pixels that were outside of the cell’s cortex). Wound edge junction intensities were normalised to the mean intensity of the 10 junctions in the surrounding tissue for each time point. The relative normalised intensity of wound edge junctions was then calculated by dividing by the mean normalised intensity of wound edge junctions in the time point prior to wounding (t0). A LOWESS curve was fitted to the data using Prism (GraphPad).

### Assigning cell row fates

Cells were assigned row identities using the “CELL_COLOR_TAG” CellOverlay tool in Epitools^6^. This was done prior to wounding and these identities were propagated in time through the entire movie. Cells could therefore change rows over time (by intercalation), but still retained their initially assigned identities (Supplementary Video 5).

### Vertex model

We model the apical surface of the tissue as a 2D network of polygonal cells, with cell-cell interfaces represented by straight edges, and three way junctions by vertices. The total mechanical energy of the tissue is given by^7^:

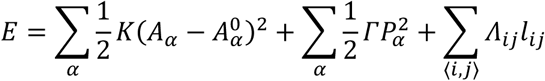

where the individual cells are labelled by *α*, and the edges connecting vertices *i* and *j* by 〈*i,j*〉 (Supplementary Fig. 2). The first term represents the area elasticity of the cells, with elastic modulus *K, A_α_* is the area of cell *α* and 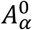 is the preferred area. The second term represents contractile energy of the actomyosin cortex for a cell with perimeter *P_α_*, and contractile tension, *Γ*. The final term represents the energy cost due to line tensions *Λ_ij_* acting on edges of length *l_ij_* which is a combination of cell-cell adhesion and cortical tension. Negative line tension implies cell-cell adhesion dominates over cortical tension, such that cells tend to maximize the length of junctions between their neighbors. The net mechanical force acting on the vertex *i* is given by ***F****_i_ = −∂E/∂****x****_i_*. Assuming over-damped dynamics, the equation of motion for vertex *i* is:

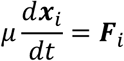

where *μ* is the coefficient of friction.

In this study, we non-dimensionalize energy by 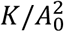 and length by 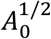, resulting in normalized contractility 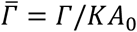 and normalized line tension 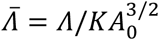. Time is non-dimensionalized by the *T* = μ/KA_0_*. The equation of motion is discretized as:

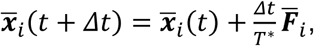

where *Δt* is the timestep. See Table 2 for a complete list of default parameter values.

**Table 2.** Default parameters used

| Parameter | Symbol | Value |
| --- | --- | --- |
| Normalized contractility | $\bar{\Gamma}$ | 0.04 |
| Normalized cell line tension | $\bar{\Lambda}_0$ | 0.00 |
| Normalized purse string tension | $\bar{\Lambda}_{ps}$ | 0.18 |
| Preferred cell area | $A^0$ | 100 $\mu\text{m}^2$ |
| Time scale | $T^*$ | 250 s |
| Cell line tension deviation | $\sigma_m$ | 0.10 |
| Tension recovery time | $\tau_m$ | 150s |
| Wound radius | $R_w$ | 1.5 |
| Time step | $\Delta t$ | 6.25 s |
| Intercalation threshold length | $L_{T1}$ | 1 $\mu\text{m}$ |
| Mean division time |  | 22.2 hours |

As the tissue relaxes by minimizing mechanical energy, cell edges may shrink due to contractile forces. If an edge length goes below a small threshold length, *L_T_*_1_, an intercalation, or T1 transition, occurs, in which a new edge is formed perpendicular to the original junction, if it results in a lower energy. To keep the system out of equilibrium, we introduce two sources of activity: cell division, and line tension fluctuations, as described below.

#### Cell division

we implemented a simplified model of the cell cycle, with cells in one of three phases; resting, growing or dividing. Cells start in the resting phase, with the default preferred area *A*_0_. Once they have reached a threshold age, they transition to mitosis at a fixed rate. In mitosis, the preferred area of the cell doubles over a period of 30 minutes, resulting in growth of the cell. After 30 minutes, the cell is divided into two new cells by creating a new edge between two of the cell edges which is chosen to minimize the system energy. This results in division of elongated cells along their short axis.

#### Line tension fluctuations

to model myosin II fluctuations, we allow the line tensions to fluctuate over time, adapting the model introduced in Curran *et al.^8^* the line tension *Λ_ij_* on the edge joining vertices *i* and *j* evolves over time as:

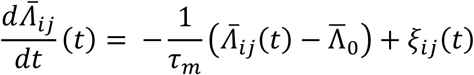

where *τ_m_* is a persistence time of myosin, 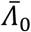 is the mean line tension, and *ξ_ij_* is an uncorrelated white noise obeying:

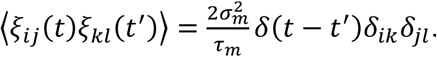

The line tension deviation, *σ_m_*, controls the amount of fluctuation around the mean, and is equal to the standard deviation of the line tension over time.

### Modelling laser ablation and wound healing

Wounds in the epithelium are created by removing any cell that lies partially or fully within a circle of radius *R_w_*. As material remains within the wound, but the actomyosin cortices are disrupted, we remove all contractility within the wound. The polygons constituting the wound have an area elasticity term with elastic modulus *K_w_* and zero contractility. As tissue material leaves the gap during wound closure, the elastic modulus decreases to zero over 10 minutes. At the same time, tension in the purse-string surrounding the wound increases from the mean line tension, 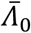, to 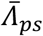, the purse-string tension. The resulting effect is a rapid expansion of wound area after ablation, followed by a contraction back to the original size over 10 minutes. We simulate the tissue dynamics for 300 minutes, or until the wound closes.

### Model implementation

The model is implemented using Surface Evolver^9^. 200 cells are generated using a Voronoi tessellation and relaxed, without fluctuations and divisions, to a steady state. The simulation is then run, with divisions and fluctuating line tensions, until the tissue has grown to 250 cells. Next, the wound is ablated, and the simulation is run until wound closure or for a maximum of 300 minutes.

### Model parameters

We use the same normalized contractility 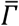 as in Farhadifar *et al.^7^*. Due to differences between our models, such as division rules, the same set of parameters do not apply. The mean division time is taken from Heller *et al.^6^*. The wound radius is chosen to give a wound with a similar number of initial edges as in experiments. We use the same tension recovery time for line tension *τ_m_* and time scale *T\** as in Curran *et al.^8^*. The line tension deviation *σ_m_*, normalized cell tension 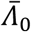, and normalized purse-string tension 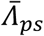 are fit against wounded and unwounded intercalation rates, and wound closure time in WT, *Mbs* RNAi, and *Rok* RNAi experiments.

### Live imaging of embryos

Adult *yw*; Ecad-tdTomato flies were placed in laying cages sealed by agar plates (water 70% v/v, apple juice 30% v/v, agar (Sigma) 3% w/v, methylparaben (Sigma) 0.05% w/v) coated by a small amount of yeast overnight at 25°C. Embryos were recovered by rinsing the agar plates with water into a basket. Embryos were dechorionated for ~1 minute in 12% sodium hypochlorite solution and rinsed thoroughly with water. Embryos were returned to agar to prevent desiccation and stage 13/14 embryos were selected by eye. Stage 13/14 embryos were affixed to coverslips using heptane glue, with their ventrolateral sides facing the coverslip. Coverslips were then attached to metal slide frames (Leica) using double sided tape and embryos were covered in halocarbon oil 27 (Sigma). Embryos were wounded as above and imaged the same as wing discs, except that 2 minute time intervals were used after the first 25 time points had been acquired. Embryos were allowed to develop to hatching after imaging, to confirm that the imaging process had not been phototoxic. Segmentation of cells was performed as described above.

### Comparing intercalation in embryos and wing discs

As a measure of how much wound edge intercalation had occurred during embryonic and wing disc wound closure, the percentage of cells remaining close to the centre of the wound was quantified. A circle 5% the area of the original wound was drawn where the centre of the wound was immediately prior to closure. Any cell that intersected this circle immediately after wound closure was scored as having not intercalated away from the wound. The number of cells was then expressed as a percentage of the total number of starting wound edge cells.

### Single junction ablations

Nanoablation of single junctions was performed to provide a measure of junctional tension. Wing discs were mounted as for wounding experiments and imaged using the same microscope. Narrow rectangular ROIs were drawn across the centre of single junctions and this region was ablated using the same settings used to wound wing discs (see above). Wing discs were imaged continuously in a single plane using identical settings as described above, except that 10x zoom was used. The initial recoil rate of vertices at the ends of ablated junctions was quantified by measuring the change in distance between the vertices and dividing by the initial time step.

### Statistical Information

The results of all statistical test are thoroughly reported in figure legends. Appropriate statistical tests were chosen based on data distributions. Kolmogorov-Smirnov and *t*-tests were two-tailed. For biological experiments, replicates represent wing imaginal discs from different animals. Because the segmentation and tracking process is extremely labour intensive, for each genetic condition 5 replicates were used. This was deemed sufficient, as clear statistical differences could be observed between genotypes. 12 replicate vertex model simulations were run for each parameter set.

### Data Availability Statement

The data that support the findings of this study are available from the corresponding author upon reasonable request.

---

**Supplementary Figure 1.**
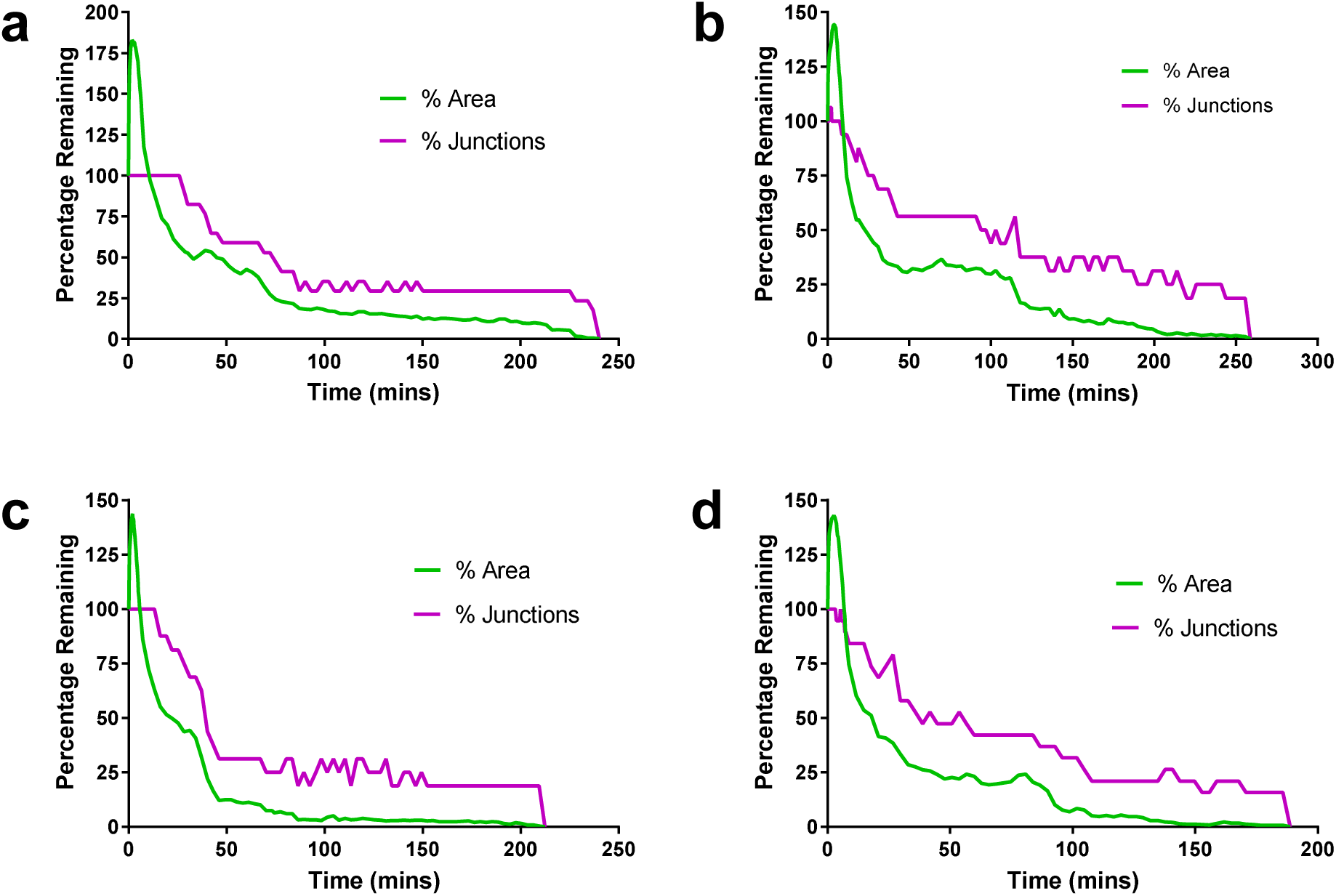
Relationship between wound area changes and intercalation events in WT wing discs. **a-d**, Quantification of the percentage of starting wound edge junctions (magenta) and wound percentage area (green) for four Ecad-GFP wing disc wounds (in addition to the wing disc in Fig. 1e).

**Supplementary Figure 2.**
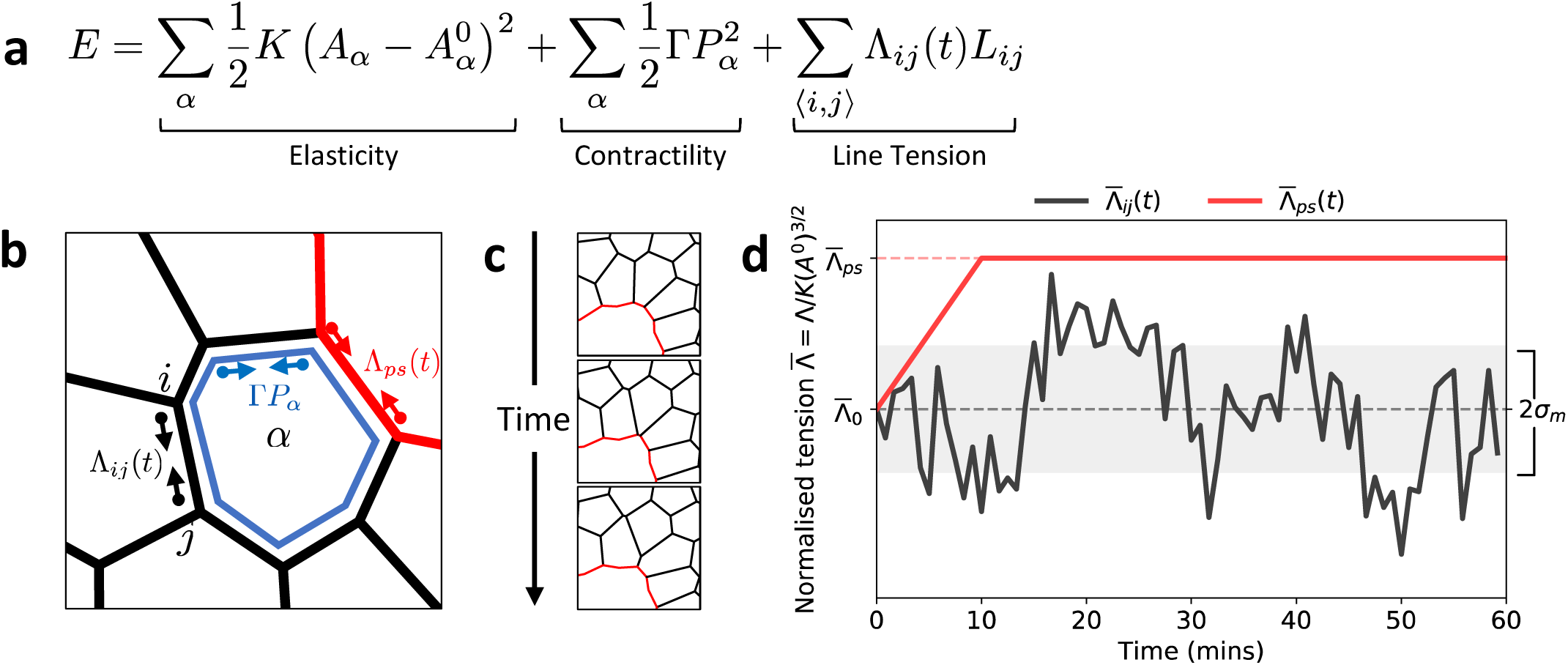
Vertex modelling of wound healing. **a**, Equation for the total energy of the system. **b**, Schematic showing the forces due to line tension and contractility acting on a cell in the vertex model. **c**, Example of a wound edge intercalation. **d**, Cell edge line tension fluctuation (black) and purse-string tension on the wound (red) over time after ablation (T = 0 mins).

**Supplementary Figure 3.**
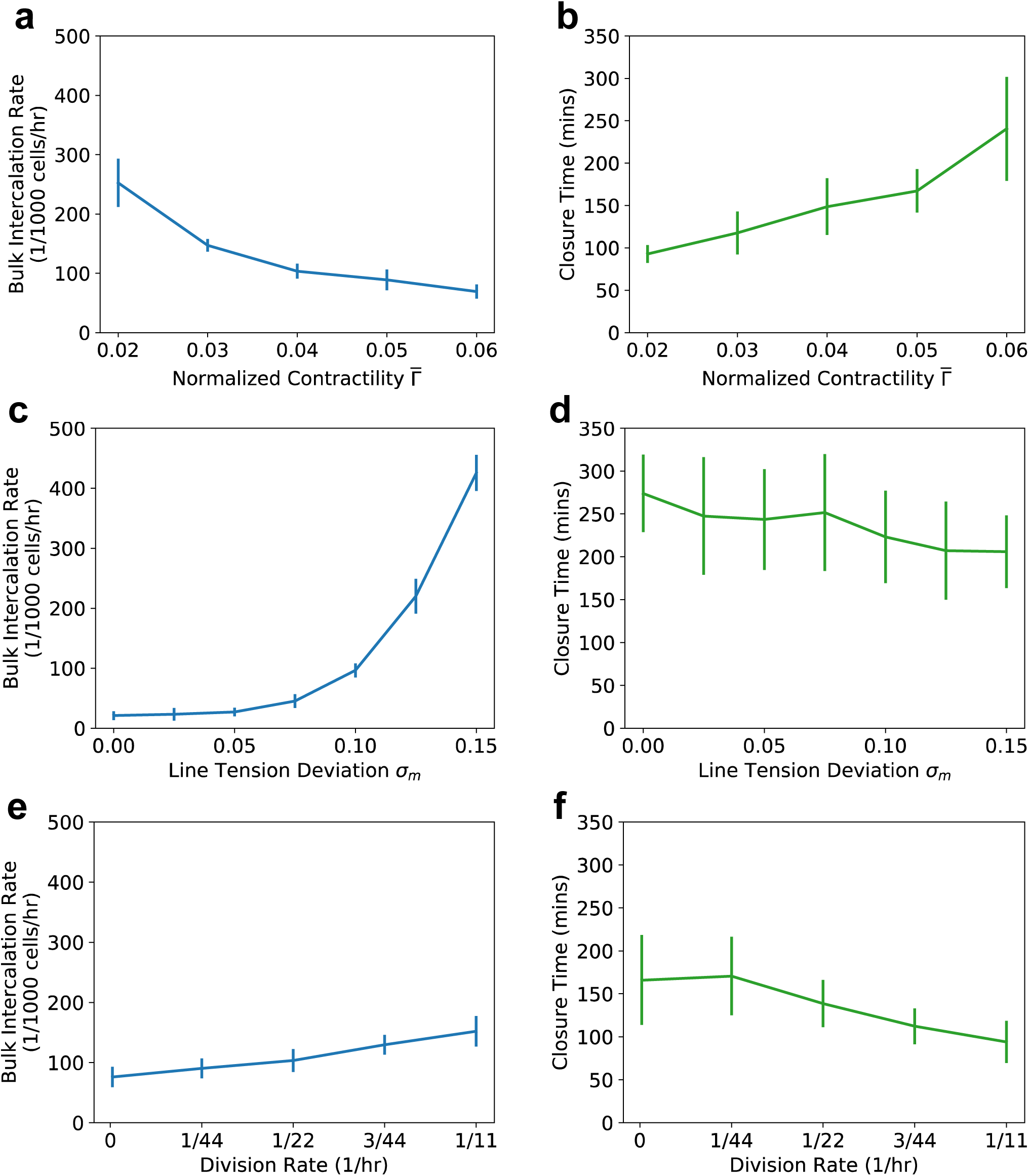
Effect of model parameters on fluidity and closure time. **a**, Mean intercalation rate in a wounded tissue, and **b**, mean closure time against normalized contractility. **c**, Mean intercalation rate in a wounded tissue, and **d**, mean closure time against cell line tension deviation. **e**, Mean intercalation rate in a wounded tissue, and **f**, mean closure time against cell division rate. For each value, (n = 12).

**Supplementary Figure 4.**
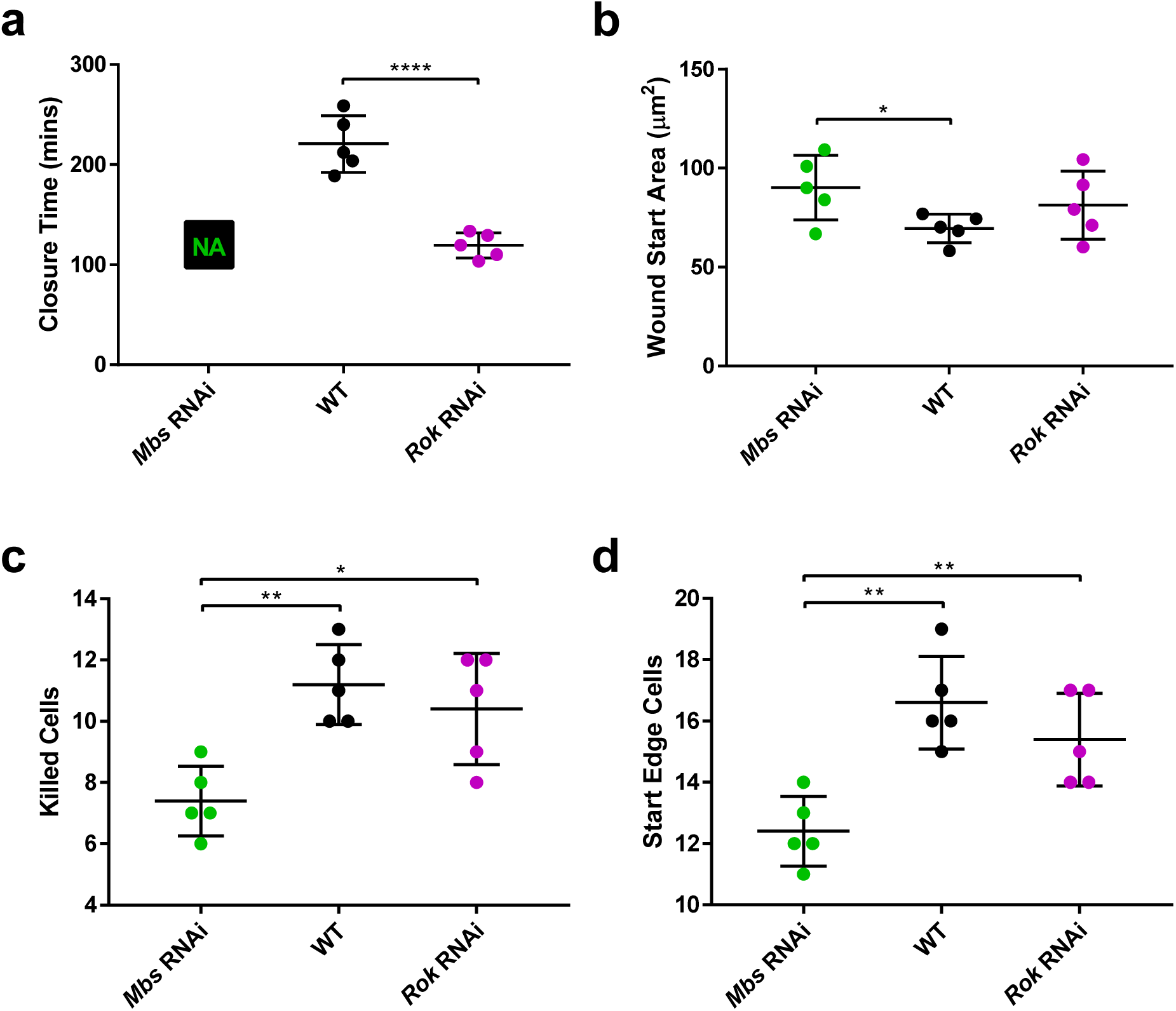
Comparisons of features of *Mbs* RNAi, *Rok* RNAi and WT wounds. **a**, Quantification of time of closure. *Rok* RNAi wounds close in less time than WT wounds (unpaired *t*-test, *t*=7.318, *df*=8, *p*<0.0001). *Mbs* RNAi wounds fail to close. **b**, Quantification of wound start areas. WT wound were smaller than *Mbs* RNAi wounds (unpaired *t*-test, *t*=2.585, *df*=8, *p*=0.0324). **c**, Quantification of number of cells killed. Fewer cells were killed in *Mbs* RNAi wounds than in WT (unpaired *t*-test, *t*=4.906, *df*=8, *p*=0.0012) and *Rok* RNAi wounds (unpaired *t*-test, *t*=3.128, *df*=8, p=0.0141). **d**, Quantification of starting wound edge cells. *Mbs* RNAi wounds had fewer starting edge cells than WT (unpaired *t*-test, *t*=4.95, *df*=8, *p*=0.0011) and *Rok* RNAi wounds (unpaired *t*-test, *t*=3.536, *df*=8, *p*=0.0077). **a-d**, Error bars = SD.

**Supplementary Figure 5.**
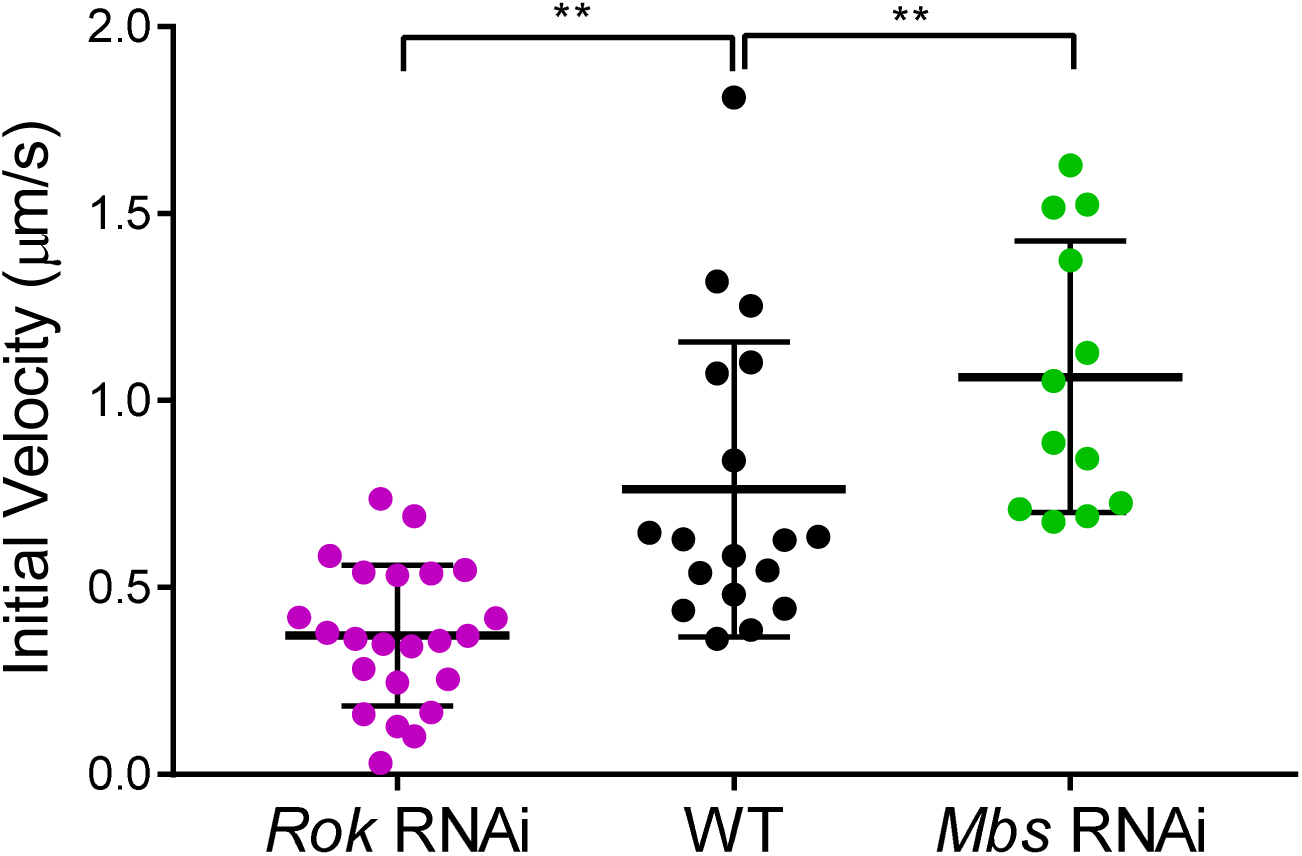
Quantifying junction tension in *Mbs* RNAi, *Rok* RNAi and WT tissues. Quantification of initial vertex recoil rates after single junction ablations in WT (black), *Mbs* RNAi (green) and *Rok* RNAi (magenta) wing discs. Compared to WT, the initial vertex recoil rate was significantly lower in *Rok* RNAi wing discs (Kolmogorov-Smirnov Test, D=0.5845, *p*=0.002) and significantly higher in *Mbs* RNAi wing discs (Kolmogorov-Smirnov Test, D=0.6667, *p*=0.0033).

**Supplementary Figure 6.**
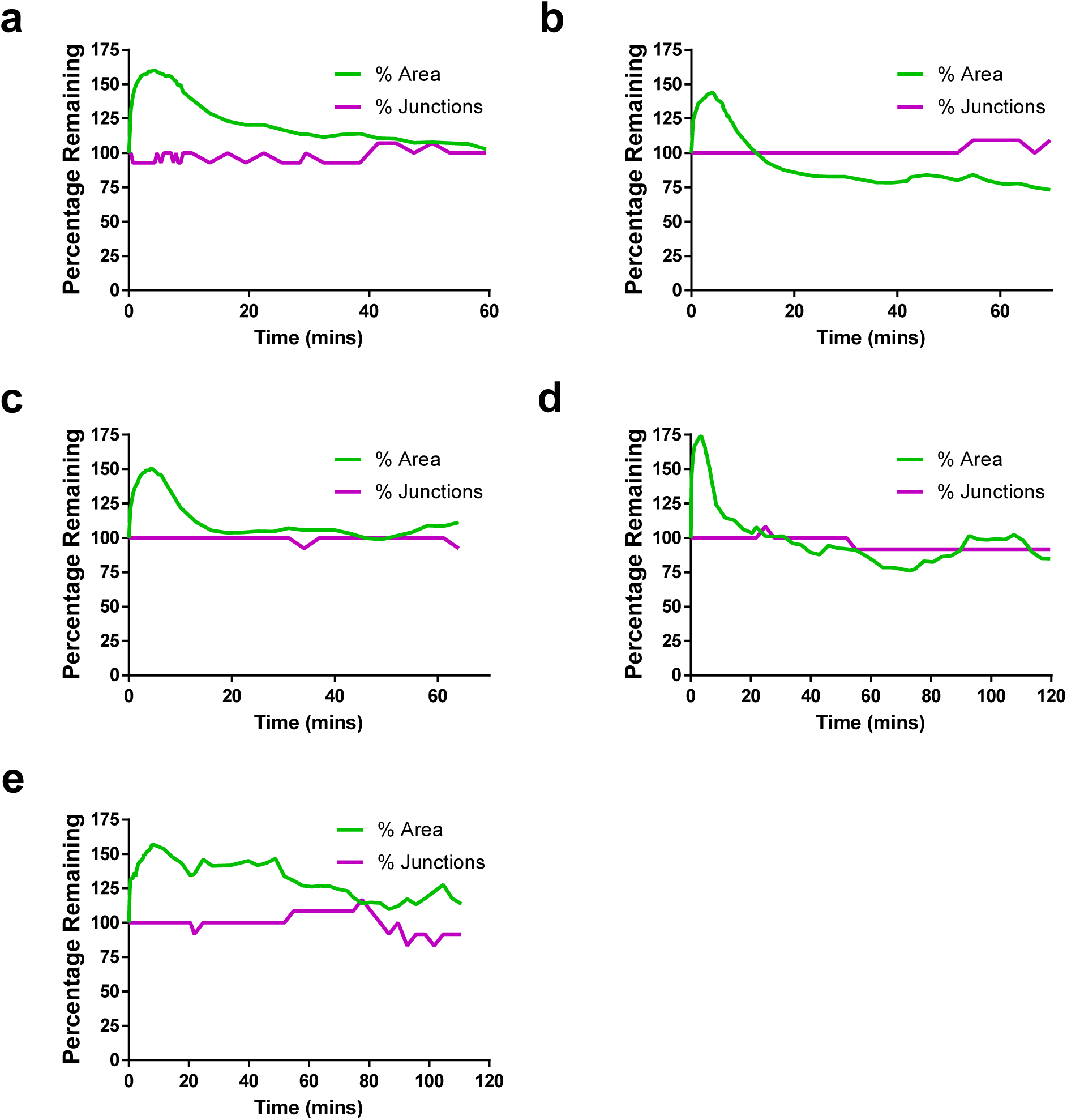
Relationship between wound area changes and intercalation events in *Mbs* RNAi wing discs. **a-e**, Quantification of the percentage of starting wound edge junctions (magenta) and wound percentage area (green) for five Ecad-GFP/UAS-*Mbs*-RNAi; *rn*-GAL4/+ wing disc wounds.

**Supplementary Figure 7.**
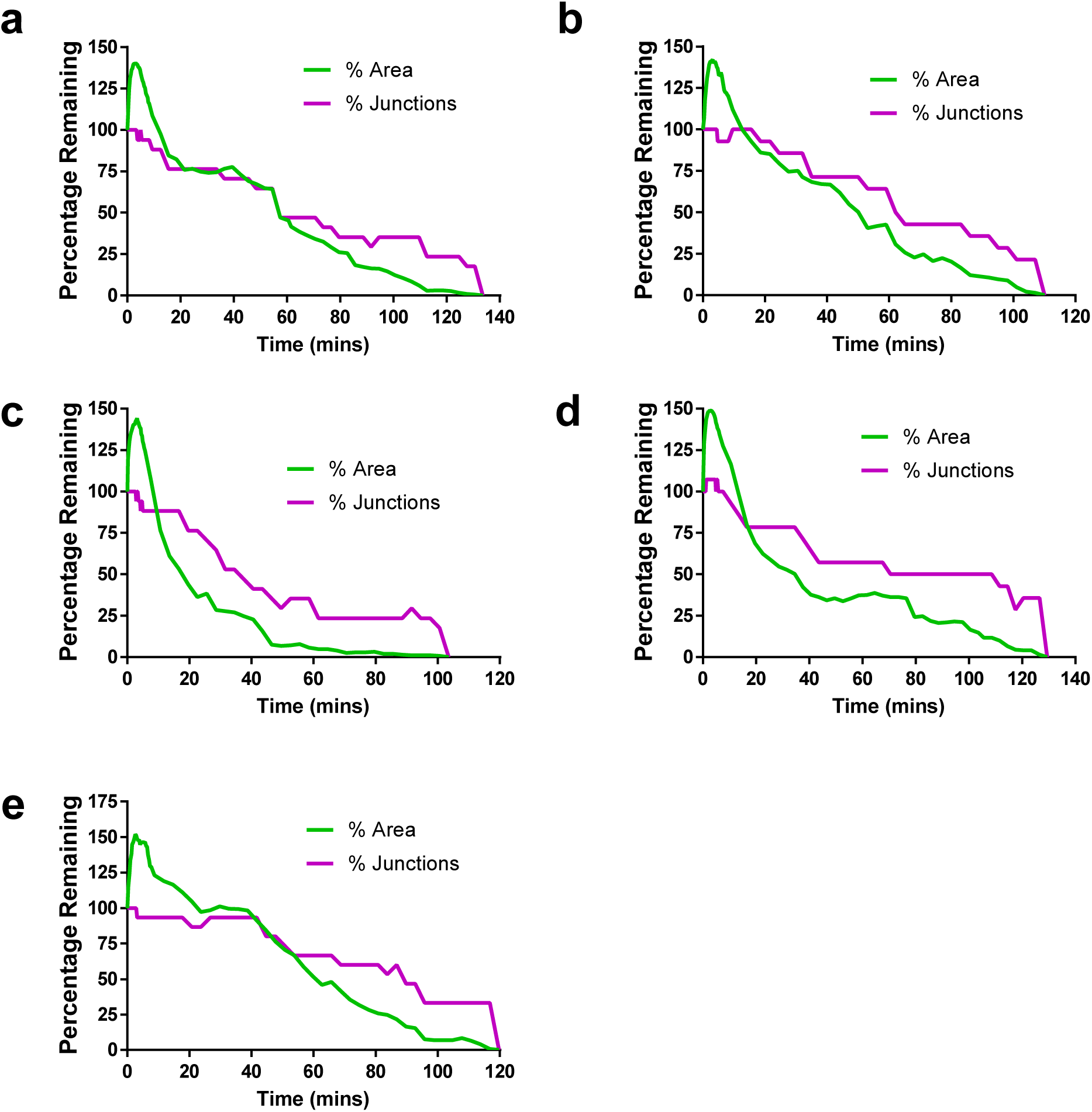
Relationship between wound area changes and intercalation events in *Rok* RNAi wing discs. **a-e**, Quantification of the percentage of starting wound edge junctions (magenta) and wound percentage area (green) for five Ecad-GFP/UAS-*Rok*-RNAi; *rn*-GAL4/+ wing disc wounds.

**Supplementary Figure 8.**
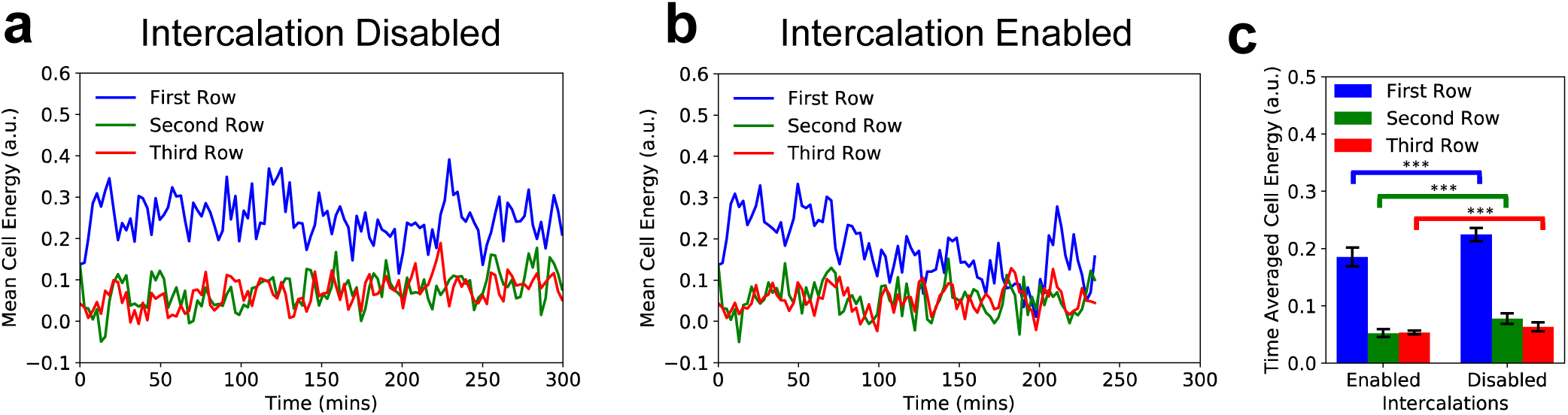
Effect of intercalations on cell energy. **a-b**, Mean cell mechanical energy over time for the first three rows of cells with intercalations (**a)** enabled, and (**b)** disabled. **c**, Quantification of time averaged cell energy for the first three rows, with intercalations enabled and disabled, averaged over 12 simulations. With intercalations enabled, the mean cell area is lower in the first row (unpaired *t*-test, *t*=6.77, *df*=22, *p*<1e-5), second row (unpaired *t*-test, *t*=7.398, *df*=22, *p*<1e-6), and third row (unpaired *t*-test, *t*=3.819, *df*=22, *p*<1e-3). Error bars = SD.

**Supplementary Video 1. Myosin II localisation during *Drosophila* wing disc wound closure**. Time-lapse of the first hour after wounding in a *sqh^AX^*^3^; *sqh*-GFP, Ecad-tdTomato wing imaginal disc. Myosin II is marked by Sqh-GFP (magenta, maximum intensity projection) and cell outlines by Ecad-tdTomato (green, adaptive projection). Myosin II rapidly accumulates at the wound’s edge in the manner of a purse string.

**Supplementary Video 2. WT wound closure**. Time-lapse of an Ecad-GFP/+; *rn*-GAL4/+ (WT) wing disc from wounding to wound closure. An adaptive projection of Ecad-GFP (greyscale) is overlaid with skeletonised cell outlines (cyan). The wound itself (white) and initial wound edge cells (blue) are highlighted.

**Supplementary Video 3. Vertex model simulation of wound healing with intercalations disabled**. Red edges represent the wound edge and have increased line tension compared to the surrounding tissue (black edges). The wound fails to close during the simulation.

**Supplementary Video 4. Vertex model simulation of wound healing with intercalations enabled**. The same parameters are used as in Supplementary Video 3, except that intercalations are enabled. The wound is able to close.

**Supplementary Video 5. Analysing the first three rows of cells away from the wound**. Time-lapse of an Ecad-GFP/+; *rn*-GAL4/+ (WT) wing disc from wounding to wound closure. An adaptive projection of Ecad-GFP (greyscale) is overlaid with skeletonised cell outlines (red). The wound itself is highlighted in white. The first (blue), second (green) and third (red) rows of cells away from the wound are selected prior to wounding. These initial cell identities are propagated through time, regardless of whether the cell intercalates or not.

**Supplementary Video 6. Wound closure in an *Mbs* RNAi wing disc**. Time-lapse of an Ecad-GFP/UAS-*Mbs*-RNAi; *rn*-GAL4/+ wing disc after wounding. The wound fails to close during imaging. An adaptive projection of Ecad-GFP (greyscale) is overlaid with skeletonised cell outlines (cyan). The wound itself (white) and initial wound edge cells that do intercalate (blue) and do not intercalate (cyan) are highlighted.

**Supplementary Video 7. Wound closure in a *Rok* RNAi wing disc**. Time-lapse of an Ecad-GFP/UAS-Rok-RNAi; *rn*-GAL4/+ wing disc from wounding to wound closure. An adaptive projection of Ecad-GFP (greyscale) is overlaid with skeletonised cell outlines (cyan). The wound itself (white) and initial wound edge cells (blue) are highlighted.

